# Charge-driven remodelling of monomeric Tau determines its phase behaviour

**DOI:** 10.64898/2026.09.21.752903

**Authors:** Yudisthira Oktaviandie, Zexiang Han, Katherine Stott, Tuomas P.J. Knowles

## Abstract

Tau transitions from a functional microtubule-stabilising monomer into pathological aggregates in tauopathies. However, the molecular mechanism by which soluble Tau transitions into insoluble aggregates remains poorly understood. Here, we show that modulation of Tau’s internal charge-charge interactions by pH and ionic strength is sufficient to remodel its monomeric conformational ensemble and direct its subsequent self-assembly pathway. By subjecting the expanded random coil-like 2N4R Tau to prolonged incubation at low ionic strength and near its isoelectric point, we generated a long-lived compact molten globule-like monomer. Although both expanded and compacted conformations remain disordered, they differ markedly in their local solvent accessibility and dynamics. We show that monomer compaction is stabilised by weak interactions that remodel the solvent accessibility of the microtubule-binding repeat region and its flanking sequences. Consequently, these conformational changes alter Tau phase behaviour, inhibiting liquid-liquid phase separation and amyloid fibril formation. Our findings demonstrate that charge-driven remodelling of the Tau conformational ensemble determines its phase behaviour and provide a mechanistic insight into the molecular events underlying tauopathies.

## Introduction

Tau’s disordered properties play essential roles in neuronal function^1,2^. However, Tau can also assemble into distinct neurofibrillary tangles (NFTs) that exhibit a highly consistent ultrastructure within a given disease and across individual patients^3–8^. Accumulating evidence has suggested that conformational rearrangements of monomeric Tau in solution are an early key event in tauopathies^9–12^. Despite our current understanding of Tau and tauopathies, the molecular mechanism that mediates soluble Tau transitions into aggregation-prone conformations remains poorly understood.

Growing evidence indicates that post-translational modifications are important in shaping the Tau conformational energy landscape. In Alzheimer’s disease, pathological Tau progressively accumulates a range of abnormal post-translational modifications (PTMs), including phosphorylation, ubiquitination, and acetylation^13^. These PTMs have been proposed to modulate Tau via direct interactions with the backbone (for phosphorylation), or via electrostatic modulation^14–16^. A combination of serine and threonine phosphorylation, as well as lysine acetylation and ubiquitination, may induce local neutralisation of positive charge that reshapes the Tau conformational landscape. Alternatively, PTMs have also been proposed to alter the cellular interactome, promoting cellular aggregation^17,18^.

Here, we investigated the mechanism of monomeric Tau compaction through charge modulation and its consequences for its phase behaviour. We were able to generate a long-lived, molten globule-like, compacted monomeric state by subjecting random coil-like, expanded Tau to prolonged incubation under low ionic strength near its isoelectric point. A combination of biophysical techniques and NMR spectroscopy confirmed that both expanded and compacted Tau remained monomeric and disordered while adopting distinct global conformational ensembles. We also demonstrated that the compacted state is stabilised by weak, non-cooperative interactions mainly located within the microtubule-binding repeat region (MTBR) and its flanking sequences. Moreover, the two conformations exhibited distinct propensities for liquid-liquid phase separation (LLPS) and amyloid fibril formation, with expanded Tau displaying a markedly greater propensity for both processes than compacted Tau. Collectively, our findings demonstrate that local charge modulation is sufficient to reshape the conformational ensemble of soluble Tau and consequently biases its subsequent phase behaviour.

## Results

### 2N4R Tau can adopt discrete long-lived disordered ensembles with distinct characteristics

To investigate how monomer conformation influences Tau phase behaviour, we first established a reproducible method to generate a long-lived compact conformational state. Our approach is summarised in Fig. 1a. Tau compaction, as assessed by microfluidic diffusional sizing (MDS), was achieved by prolonged dialysis of expanded Tau under dilute conditions and low ionic strength (10 mM) near the isoelectric point (pI ∼ 8.8). We chose this condition to minimise aggregation while promoting long-range interactions and mimicking the net charge shift as induced by PTMs in disease^13^. Following the treatment, and upon returning to a physiological ionic strength (160 mM), compacted Tau exhibited a markedly different pH-dependent response compared to expanded Tau under the same condition (Fig. 1b). The treated Tau adopted its most compact conformation near its isoelectric point, expanding at both lower and higher pH, while the untreated Tau appeared to be stably expanded, insensitive to changes in pH value. The estimated hydrodynamic radii (R_h_) for two theoretical coils, random coil and molten globule (ν = 0.5 and 0.33, respectively), are shown for comparison purposes.

**Figure 1.**
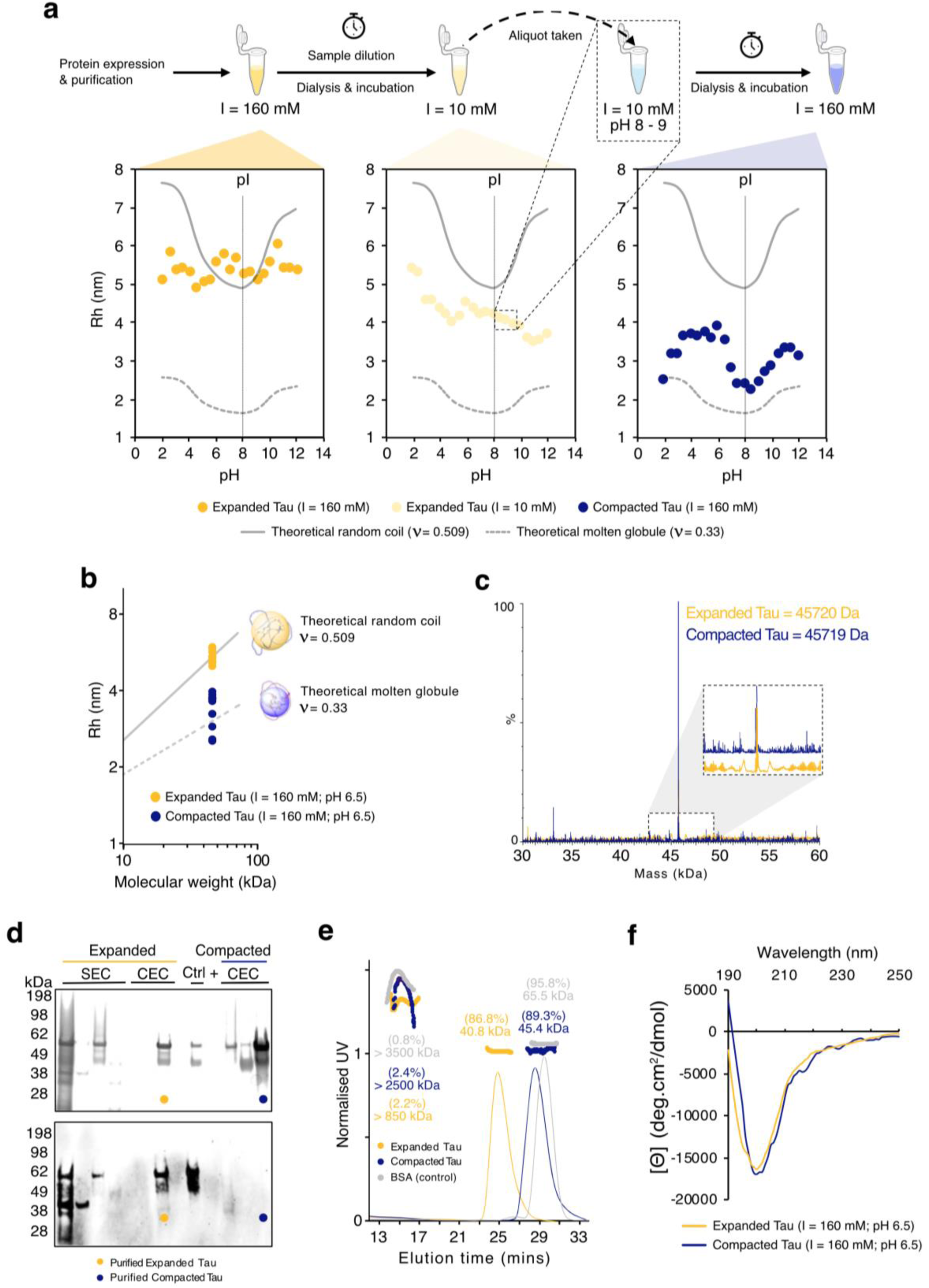
**a,** Schematic diagram of sample preparation used in this study and the hydrodynamic radii (R_h_) of the samples in Britton-Robinson (BR) universal pH buffer system^19^ at varying pH. Expanded Tau (yellow) was pre-treated in low ionic strength buffer (pale yellow) to induce compaction producing compacted Tau (dark blue). (This nomenclature and colour coding indicating expanded Tau and compacted Tau are used throughout.) **b,** R_h_ of expanded and compacted Tau plotted against their molecular weight relative to the estimated sizes of different protein states ^20^. **c,** Mass profile of expanded and compacted Tau. **d,** Non-reducing SDS-PAGE (above) and Western Blot (below) of expanded and compacted Tau at different purification steps. **e,** Elution volume and estimated molecular weight of expanded and compacted Tau, alongside BSA (grey) as a control, detected by SEC-MALS in PBS with 5 mM TCEP at pH 7.2. **f,** CD spectra of expanded and compacted Tau.

Multiple biophysical techniques confirmed that compaction alters the global dimensions of Tau while also retaining its monomeric and disordered nature. Data collected at a single pH value (6.5) and near-physiological ionic strength (160 mM) are shown in Fig. 1b-e. Under these conditions, we confirmed the R_h_ and apparent global dimensions using MDS (Fig. 1b). The expanded and compacted conformations were confirmed to be chemically identical with no detectable modifications introduced during the compaction by mass spectrometry (Fig. 1c). Non-reducing SDS–PAGE and Western blotting demonstrated that both conformations migrated with similar apparent molecular weights and showed no evidence of high-molecular-weight (HMW) species formed by intermolecular disulphide bonds (Fig. 1d). We verified this orthogonally by utilising size exclusion chromatography coupled to multi-angle light scattering (SEC-MALS) which showed that both conformations of Tau are monomeric (Fig. 1e). Interestingly, compacted Tau generated from the same batch as expanded Tau, was not detected by Western blotting (Fig. 1d, bottom). However, in a different batch preparation, compacted Tau was detected using the same antibody, albeit with reduced band intensity (Supplementary Fig. 1). This observation suggests that compaction alters the accessibility of the antibody epitope. We were therefore surprised to observe that the circular dichroism (CD) spectroscopy of both compacted and expanded Tau displayed spectra consistent with disordered proteins without any noticeable structural differences (Fig. 1f).

### Tau monomer collapse is modulated by its charge distribution

We further investigated the role of charge in Tau compaction by replotting the R_h_ data from Fig. 1b vs. theoretical net charge. Compacted Tau demonstrated a unique sensitivity to pH: as the net charge approaches zero, a steep compaction takes place, after which the R_h_ increases gradually with increasing negative charge (Fig. 2a, dark blue circles) as expected from a charged molten globule-like chain (ν = 0.33) (Fig. 2a, dashed grey line). Expanded Tau (Fig. 2a, yellow and pink circles) shows no such sensitivity but nevertheless deviates from the theoretical behaviour of a charged random coil-like chain (ν = 0.509) (Fig. 2a, solid grey line). This behaviour implicates a charge-dependent mechanism underlying Tau compaction, resulting in conformational changes that respond distinctively to pH perturbation.

**Figure 2.**
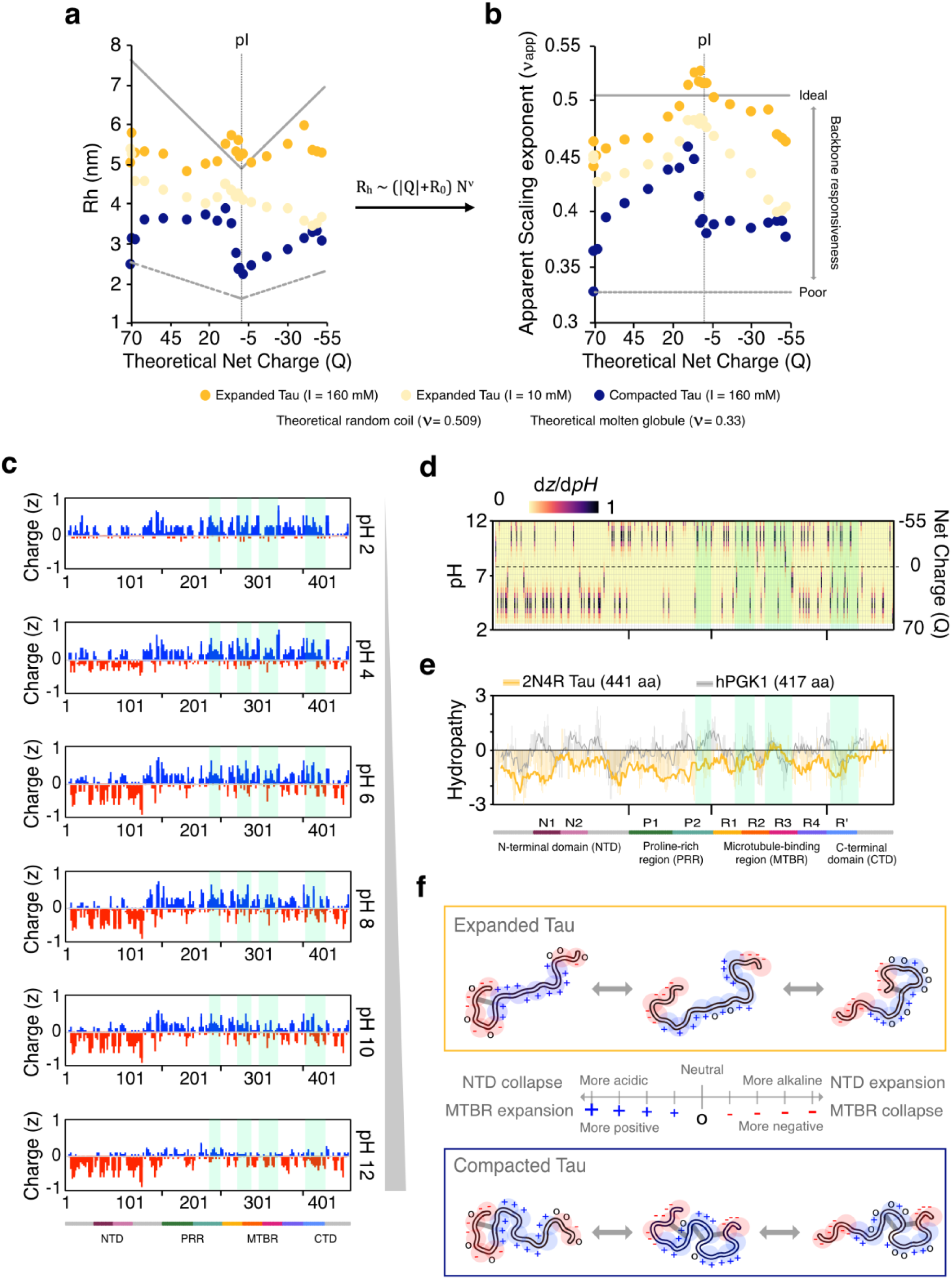
**a,** Hydrodynamic radii, and **b,** apparent Flory scaling exponent (ν) of expanded (yellow) and compacted (dark blue) Tau plotted against Tau’s theoretical net charge across a range of pH values, as indicated in Fig. 1a. **c,** Charge patterning of 2N4R Tau sequence across pH, and **d**, charge alteration in response to changes in pH (dz/dpH). **e**, Hydropathy plot of the 2N4R Tau sequence (yellow) compared with the folded human PGK1 protein (44.6 kDa; grey). Coloured boxes at the bottom indicate different segments (N1 – R’), and the faded light green box within each plot indicates segments with significant solvent accessibility changes shown in Fig. 4c. **f**, Schematic diagram of the electrostatic effect on the expanded and compacted Tau conformations.

The same data are replotted in Fig. 2b with R_h_ converted to the Flory exponent ν to highlight this effect. The size of a polyelectrolyte is generally expected to be a minimum as the net charge approaches zero^21^. However, as the net charge approached zero, the blocky charge patterning of Tau (Fig. 2c) resulted in electrostatic repulsion between like-charged regions that promoted swelling, as evidenced by expanded Tau in either salt condition (Fig. 2b). Moreover, it also demonstrated the effect of ionic strength, indicating general relative compaction (vertical displacement). Similar behaviour was seen for compacted Tau in the positive net charge regime, while it appeared to be conformationally ‘inert’ above the pI, as indicated by lack of ν response to the changing overall net charge (Fig. 2b).

The conformational differences and distinct pH responses between expanded and compacted Tau may be reflected in their charge distribution vs. pH (Fig. 2c). Overall, the Tau N-terminal and C-terminal domains (NTD and CTD, respectively) are enriched with negatively charged residues, while the proline-rich and microtubule-binding regions (PRR and MTBR, respectively) are highly positively charged. Titration below the isoelectric point mainly affects the charge within NTD and CTD, while the PRR and MTBR charge is modulated mainly above the isoelectric point (Fig. 2c). To highlight these changes across the sequence, a plot showing the first derivative is also shown (Fig. 2d). It reveals a more concentrated presence of titratable residues (histidine and cysteine residues, which have intermediate pKa values closer to neutral pH) within the MTBR and the pseudo-repeat R’ in the CTD. Together with the observations shown in Fig. 2a & 2b, this suggests that Tau compaction is driven by changes that are predominantly in its PRR and MTBR. Given the possible involvement of cysteine residues, the compaction by disulphide bond formation was excluded by titration with a strong reducing agent (Supplementary Fig. 2a). No change in R_h_ was observed upon reduction, demonstrating that the compacted state is not stabilised by disulphide bonds. Further inspection of Tau’s hydropathy (Fig. 2f) reveals that residues located within the MTBR and CTD are among the most hydrophobic, suggesting that Tau compaction is potentially mediated by non-covalent interactions in these regions.

A denaturant titration indicated some expansion of compacted Tau, while the expanded Tau remained relatively unchanged (Supplementary Fig. 2b). Notably, addition of 6 M Gdm-HCl did not disrupt the compaction completely, supported by the previous observation of several 8 M-Urea-resistant turn conformations within MTBR^22^. The continuous, non-cooperative expansion of compacted Tau, in contrast to the typical sigmoidal response of a folded protein reference (hPGK1)^23^, indicates that the compacted state is stabilised by local, independent interactions that are likely weak and multivalent rather than part of a cooperative folding mechanism. Moreover, anion-dependent expansion of compacted Tau (Supplementary Fig. 2c) reveals that Tau compaction can be disrupted by small, amphiphilic ions (e.g., acetate & succinate), suggesting a combination of both electrostatic and hydrophobic interactions in mediating compaction.

The effect of local charge perturbation on Tau conformation is summarised in Fig. 2f. The conformational space of compacted Tau is distinct from expanded Tau due to MTBR re-organisation stabilised by non-covalent interactions. In expanded Tau, the negatively charged NTD collapse under acidic conditions is compensated by positively charged MTBR extension, and vice versa in an alkaline environment. Meanwhile, because MTBR is restricted in compacted Tau, it can only be partially extended at low pH below the isoelectric point and appears unresponsive at high pH above the isoelectric point.

### Tau compaction involves a dynamic re-arrangement of the backbone within MTBR and its flanking regions affecting its solvent interactions

To further understand the molecular basis of these Tau conformations, we employed NMR spectroscopy to probe distinct characteristics of protein dynamics and solvent exposure. ^1^H-^15^N-HSQC measurements (Fig. 3a) of the expanded and compacted Tau are similar and closely resemble previously assigned spectra of 2N4R Tau, allowing us to transfer ∼88% of the assignments with confident^24^. Although both spectra overlap reasonably well and show the typical ^1^H^N^ chemical shift dispersion of a disordered protein, the peak intensities were not uniform across the sequence, consistent with local secondary structure and long-range interactions observed previously^25,26^. Expanded Tau showed reduced intensities predominantly in the PRR, while compacted Tau showed widely-distributed reduced intensities from the PRR to the C-terminus. Interestingly, a similar stretch of residues was shown previously to have elevated R1**⍴** relaxation rates, indicating slower motions^26^. Moreover, close inspection revealed contiguous regions with small chemical shift differences between the expanded and compacted Tau (Fig. 3b) concentrated around histidine residues and more generally in the MTBR and CTD (Supplementary Fig. 3). Given that the buffer conditions of both samples were identical and achieved through extensive dialysis, the small displacement of histidine chemical shifts may indicate a local pK_a_ shift due to conformational rearrangement^27,28^. Interestingly, the peak intensity of residues at the P1-P2 boundary, which was already relatively weak compared with other regions in both conformations, remained largely unchanged upon compaction (ΔIntensity; Fig 3a, bottom). This observation suggests that the apparent global reduction in peak intensity in the compacted state is unlikely to result from differences in protein concentration or data acquisition. Instead, the widespread peak broadening in compacted Tau indicates altered conformational dynamics compared to expanded Tau, potentially arising from intermediate exchange on the microsecond-to-millisecond timescale^29^.

**Figure 3.**
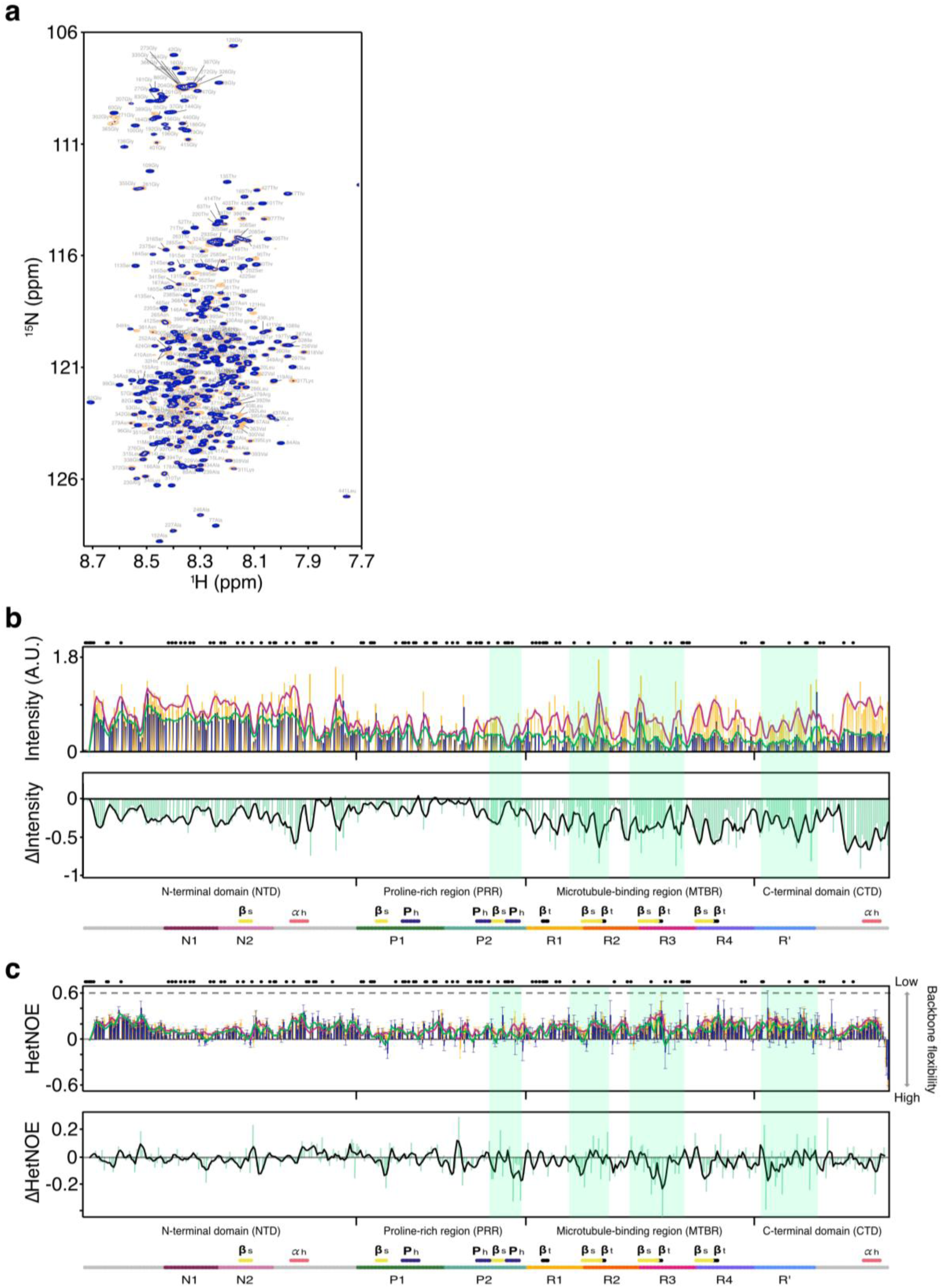
**a**, ^1^H-^15^N-HSQC NMR spectra of expanded (yellow) and compacted (dark blue) Tau in 10 mM sodium phosphate, 150 mM, 3 mM NaN_3_, 5 mM DTT, 10% D_2_O pH 6.8. **b**, Peak intensity (top) of expanded (yellow bars) and compacted (dark blue bars) and their interpolated trendline shown as magenta and green lines, respectively. Peak intensity differences (ΔIntensity; bottom) (light green bar) and its interpolated trendline shown as black lines. **c**, HetNOE value (top) of expanded (yellow bars) and compacted (dark blue bars) and their interpolated trendline shown as magenta and green lines, respectively. HetNOE value differences (ΔHetNOE; bottom) (light green bar) and its interpolated trendline shown as black lines. Black dots at the top, and coloured boxes at the bottom in **b** and **c** indicate unassigned residues, and different segments (N1 – R’) as well as transient structures (α_h_ = α-helix; β_s_ = β-sheet; P_h_ = polyproline-helix II; β_t_ = β-turn) as previously reported ^22,26^. Faded light green box in **b** and **c** indicate segments with significant solvent accessibility change shown in Fig. 4c. CSP, fitted *k*_ex_, fitting RMSE and ΔIntensity at t_mixing_ = 78.8 ms are shown in Supplementary Fig. 3.

Backbone dynamics measured by heteronuclear NOE (hetNOE) further demonstrated that both conformations remained globally disordered while exhibiting subtle local differences. The hetNOE of expanded and compacted Tau revealed highly similar profiles across the sequence (Fig. 3c, top), indicating that both conformations exhibit comparable fast backbone dynamics. Importantly, the hetNOE values for both states were below 0.4 throughout the sequence, consistent with the absence of persistent secondary structure characteristic of folded proteins^30^. These observations are supported by the circular dichroism data (Fig. 1d) and further confirm that both conformations remain intrinsically disordered. Despite this general similarity, closer examination revealed subtle reductions in hetNOE values upon compaction, as shown in a difference plot (ΔhetNOE; Fig. 3c, bottom), suggesting increased local flexibility occurring on the picosecond-to-nanosecond timescale despite the overall chain compaction. Notably, these regions coincide with segments previously reported to adopt transient secondary structural elements^22,26^.

Solvent exchange rates were measured using CLEANEX-PM, which indicated that residues within the MTBR and its flanking regions become less exposed to the solvent upon compaction (Fig. 4a, b, and c). Water-to-amide proton transfer was measured at multiple short mixing times and fitted to obtain residue-specific solvent exchange rates (*k*_ex_; Fig. 4b)^29,31^. Comparison of the fitted exchange rates with the corresponding intrinsic exchange rates (*k*_int_) generated residue-specific maps of relative solvent accessibility (*k*_ex_/*k*_int_) for each conformation (Fig. 4c). Although both expanded and compacted Tau exhibited similar overall solvent accessibility profiles (Fig. 4c, top), several regions displayed significant reductions in *k*_ex_/*k*_int_ value upon compaction, indicating increased backbone protection from solvent (Fig. 4c, bottom). This conclusion was supported by qualitative comparison of the CLEANEX-PM signal intensities at the longest mixing time (t_mixing_ = 78.8 ms), demonstrating similar region-specific decreases upon compaction (Fig. 4a). Moreover, in comparison to the subtle changes in fast backbone dynamics measured by hetNOE, the more substantial changes measured in this experiment further suggested that the conformational differences between Tau species are dominated by intermediate exchange.

**Figure 4.**
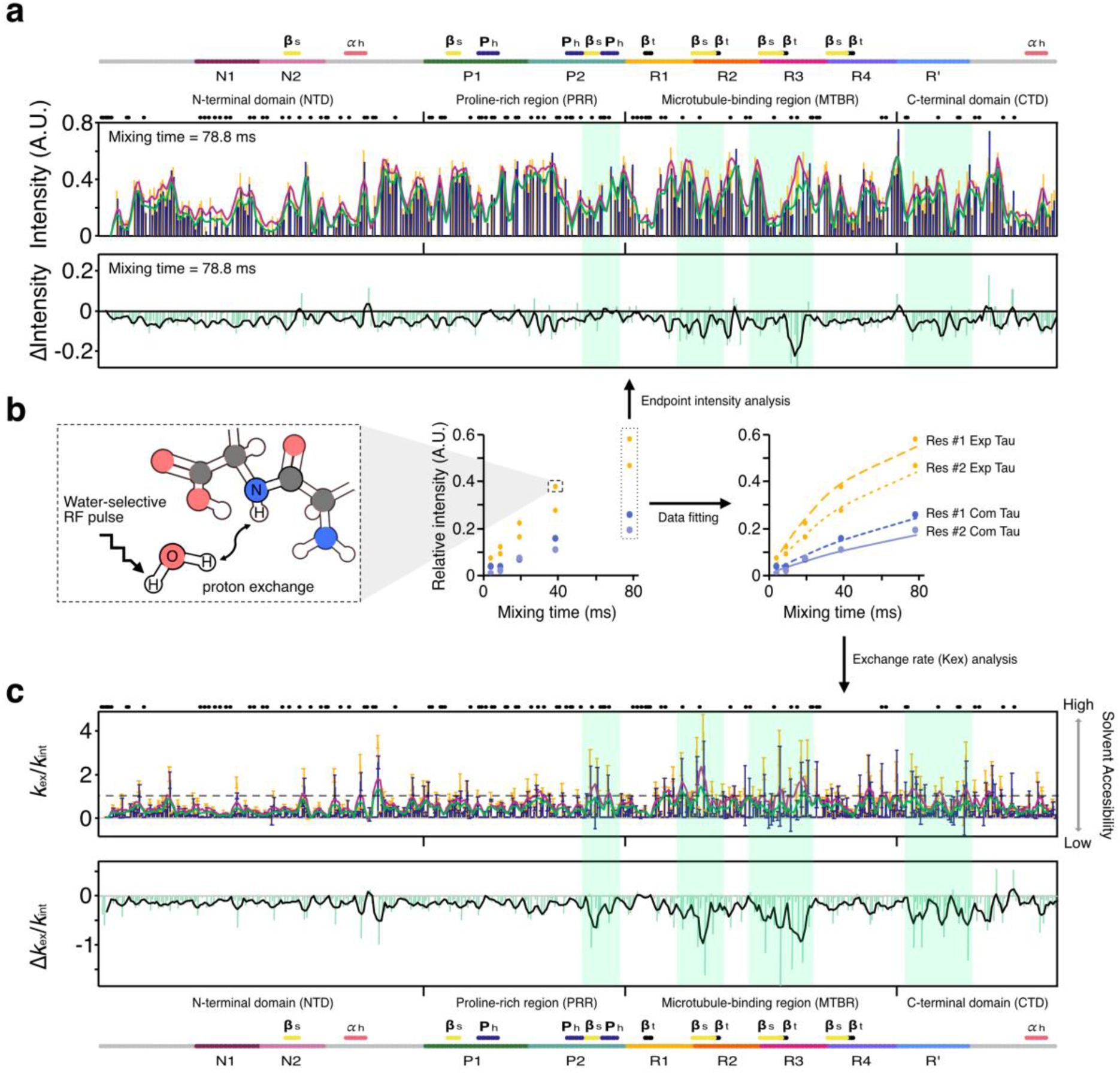
**a**, CLEANEX-PM peak intensity at t_mixing_ = 78.8 ms (top) of expanded (yellow bars) and compacted (dark blue bars) and their interpolated trendline shown as magenta and green lines, respectively. CLEANEX-PM peak intensity differences at t_mixing_ = 78.8 ms (bottom) (light green bar) and its interpolated trendline shown as black lines. **b,** Schematic diagram of magnetisation transfer from water to backbone amide and fitted residues as measured in CLEANEX-PM experiment. **c**, *k*_ex_/*k*_int_ ratio (top) of expanded (yellow bars) and compacted (dark blue bars) and their interpolated trendline shown as magenta and green lines, respectively. *k*_ex_/*k*_int_ ratio differences (Δ*k*_ex_/*k*_int_; bottom) (light green bar) and its interpolated trendline shown as black lines. Black dots at the top in **b**, **c**, and **d** indicate unassigned residues. Coloured boxes at the bottom in **b**, **c**, and **d** indicate different segments (N1 – R’) and transient structures (α_h_ = α-helix; β_s_ = β-sheet; P_h_ = polyproline-helix II; β_t_ = β-turn) as previously reported ^22,26^ Faded light green box in **b**, **c**, and **d** indicate segments with significant solvent accessibility change shown in **d**. CSP, fitted *k*_ex_, fitting RMSE and ΔIntensity at t_mixing_ = 78.8 ms are shown in Supplementary Fig. 3.

Collectively, the combined HSQC, hetNOE, and CLEANEX-PM data support a model in which expanded and compacted Tau differ primarily in their conformational dynamics rather than in their average structural architecture. Together with the denaturant and charge - response analyses, these findings support a model in which compaction is mediated by interactions involving MTBR and its flanking regions. In particular, the reduced solvent accessibility upon compaction within R3 and R’ pseudo-repeats is consistent with a potential interaction between these regions, possibly mediated by conserved IVYK hydrophobic motifs present within both regions. Importantly, despite the presence of cysteine and tyrosine residues that could potentially stabilise the collapse through covalent bond formation, the CSPs and hetNOE values for these residues argue against the formation of such cross-linking (Supplementary Fig. 4). Similarly, our NMR observations are also inconsistent with the formation of ordered higher-order assemblies, supporting the conclusion that Tau remains monomeric and disordered despite compaction.

### Exposed motifs within MTBR and its flanking regions are required for liquid – liquid phase separation and fibril formation

To understand the pathological consequences of monomeric Tau conformations, we investigated their propensity to undergo liquid – liquid phase separation (LLPS)^32^. By examining their response to temperature, we found that only expanded Tau retains the ability to form liquid-like condensates. Using the method indicated in Fig. 5a, highly concentrated expanded Tau (200 μM) displayed a spontaneous coacervation at room temperature, even without the addition of cofactors. At lower concentrations, yet remaining above the apparent saturation concentration, expanded Tau exhibited a reversible lower critical solution temperature (LCST)-type transition as indicated by two successive heating and cooling cycles (Fig. 5b). Upon addition of salt, or further dilution, the coacervates were readily dissolved (Fig. 5a, b). Moreover, although compacted Tau also exhibited apparent temperature-dependent phase separation with a higher onset temperature (Fig. 5c), it formed HMW species that progressively enriched with increasing temperature (Fig. 5d), suggesting oligomer formation rather than LLPS.

**Figure 5.**
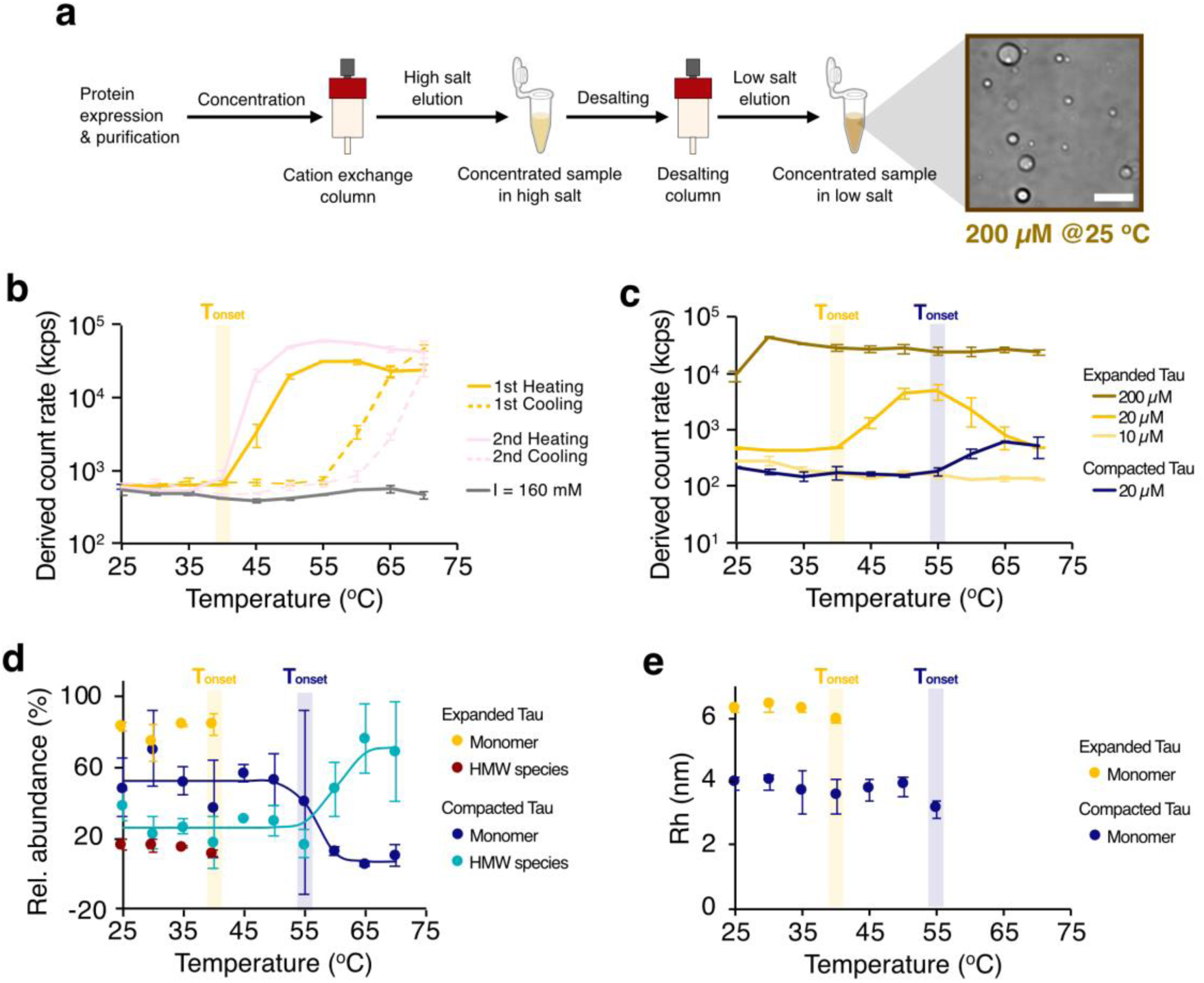
**a,** Schematic diagram of sample preparation for Tau phase separation without requiring additional cofactors. At a high concentration (200 μM), expanded Tau formed condensates that can be seen under microscope. The white bar within the micrographs indicates 15 μm. **b,** Heating–cooling cycles of self-coacervating expanded Tau in 10 mM sodium phosphate, 5 mM TCEP at pH 6 at 20 μM. Upon the addition of salt (grey), self-coacervation was not observed. **c,** Temperature-dependent phase separation of expanded (yellow) and compacted (dark blue) Tau in 10 mM sodium phosphate, 5 mM TCEP at pH 6. **d,** Relative abundance of expanded and compacted Tau monomers (yellow and dark blue, respectively) and oligomers (maroon and turquoise, respectively) in 10 mM sodium phosphate, 5 mM TCEP at pH 6 at 20 μM. **e,** Hydrodynamic radii of expanded (yellow) and compacted (dark blue) Tau below the onset temperature (faded coloured bars) in 10 mM sodium phosphate, 5 mM TCEP at pH 6 at 20 μM.

This emergence of phase-separating components above a threshold temperature is consistent with an entropy-driven phase separation process. Notably, the inability of compacted Tau to form liquid droplets further suggests that temperature-dependent phase separation requires interaction between residues within MTBR and CTD, which are more accessible in expanded Tau. Moreover, the higher onset temperature of compacted Tau indicated that this species does not undergo expansion that follows the same pathway as expanded Tau. This behaviour is indicated by the R_h_ of compacted Tau that remained unchanged above the expanded Tau onset temperature (Fig. 5e). Together, this suggests that Tau LLPS is likely associated with the release of ordered water surrounding exposed hydrophobic surfaces within the MTBR and CTD, which is supported by NMR observations (Fig. 4a, b, and c).

Furthermore, Tau–PEG phase diagrams revealed distinct charge dependencies of phase separation between expanded and compacted Tau. Using a microfluidic platform^33^ (Fig. 6a), phase diagrams generated across a range of ionic strengths demonstrated that expanded Tau exhibited an approximately symmetrical shift in phase boundaries while phase separation of compacted Tau was suppressed upon salt addition within the concentration regime examined (Fig. 6b; Supplementary Fig. 5). Characterisation of expanded Tau condensates by fluorescence recovery after photobleaching (FRAP) revealed liquid-like behaviour (Fig. 6c) with different apparent diffusion coefficient (D) across ionic strength (Fig. 6d). Strikingly, the diffusion coefficient exhibited a sigmoidal transition suggesting a two-state-like change in condensate dynamics. Together with the salt-dependent phase boundary shifts, these observations support a contribution of long-range electrostatic interactions in Tau LLPS. Importantly, the persistence of phase separation under conditions with strong electrostatic screening indicates that short-range interactions also contribute to condensate formation (Fig. 6f).

**Figure 6.**
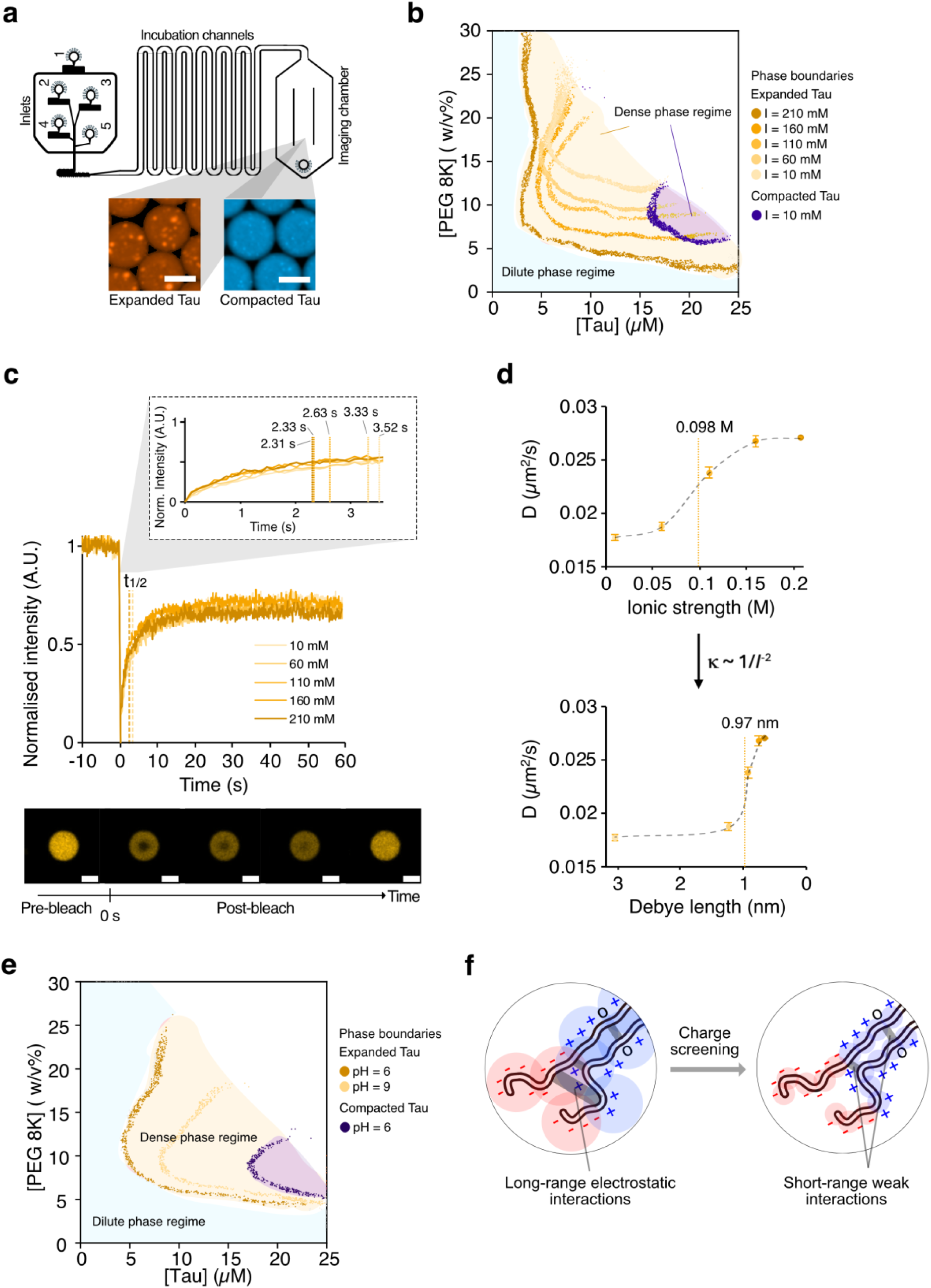
**a,** Schematic diagram of the microfluidic platform, Phasescan^33^, used to generate phase diagrams. Water-in-oil droplets containing expanded (yellow) or compacted (dark blue) Tau condensates forming puncta with known concentrations, as reported by dyes, were imaged to generate phase diagrams. The white bar within the micrographs indicates 65 μm. **b,** Phase diagram of expanded (yellow) and compacted Tau (dark blue) in 10 mM sodium phosphate, 5 mM TCEP at pH 6 with different ionic strengths. Phase boundaries separating the dilute and dense phase regimes are indicated with filled circles. **c,** Fluorescence recovery after photobleaching (FRAP) of expanded Tau condensates (20 μM Tau & 10 w/v% 8k PEG) in 10 mM sodium phosphate, 5 mM TCEP at pH 6 at varying salt concentrations. The white bars within the micrographs showing photobleaching recovery indicate 2 μm. The inlet displays the calculated fluorescence recovery half-time (t_1/2_) of Tau condensates at different ionic strengths. **d,** Calculated diffusion coefficient (D) of the fluorescent species within Tau condensates as a function of ionic strength (top) or Debye length (bottom), indicated with their respective inflection or transition points. **e,** Phase diagram of expanded (yellow) and compacted Tau (dark blue) in BR buffer at pH 6 and 9. Phase boundaries separating dilute and dense phase regimes are indicated with filled circles. **f,** Schematic diagram of charge screening effect on Tau intermolecular interaction within condensates.

The pH-dependent phase diagrams further demonstrated that Tau phase separation is governed not only by global electrostatic properties but also by local charge distribution and conformational accessibility. At pH 6, compacted Tau exhibited a markedly higher saturation concentration required for phase separation compared with expanded Tau (Fig. 6e; Supplementary Fig. 5). Under a more acidic condition (pH 4), neither expanded nor compacted Tau underwent detectable phase separation, consistent with increased intermolecular repulsion from the high excess positive net charge (Supplementary Fig. 5). Interestingly, under alkaline conditions (pH 9), where the net charge is substantially reduced, only expanded Tau retained its ability to phase separate albeit the increasing saturation concentration (Fig. 6e; Supplementary Fig. 5). This observation indicates that the local charge redistribution may selectively influence the accessibility of interaction motifs responsible for phase separation. Consistent with this conclusion, the MTBR and its flanking region, as identified by NMR (residues 224–403), experience substantial charge neutralisation at pH 9 compared with other regions (Fig. 2c). Importantly, this regional effect differs from the consequences of global electrostatic screening. While pH changes primarily shifted only the Tau concentration boundary of the phase diagram (Fig. 6e), changes in ionic strength produced more symmetrical shifts in both Tau and PEG concentration boundaries (Fig. 6b), highlighting distinct roles of local charge modulation and global electrostatic screening in regulating Tau phase separation.

Finally, aggregation assays and electron microscopy demonstrated that the conformational accessibility of the microtubule-binding repeat region (MTBR) and its flanking sequences strongly influence Tau fibril formation. Thioflavin T (ThT) aggregation assays revealed substantial differences in the total fluorescence intensity between expanded and compacted Tau (Fig. 7a). Despite this, both conformations exhibited nearly identical normalised kinetic profiles demonstrated by comparable lag times (t_lag_) and half-times (t_1/2_) (Fig. 7b). However, electron microscopy showed that expanded Tau predominantly formed fibrillar structures displaying the characteristic twisted morphology of Tau fibrils while compacted Tau generated predominantly non-fibrillar aggregates at the end of the reaction (Fig. 7c). These observations indicated that the two conformations access distinct aggregation pathways. While expanded Tau retains the ability to form assemblies that mature into ThT-positive fibrils, compacted Tau preferentially populates an alternative, disordered aggregation state. The residual ThT fluorescence observed in compacted Tau may therefore reflect a residual population of expanded Tau that remains after the compaction process. Together with previous analyses, these findings support a model in which conformational accessibility of the MTBR and its flanking regions governs soluble Tau’s ability to engage intermolecularly, biasing its phase behaviour and aggregation pathways.

**Figure 7.**
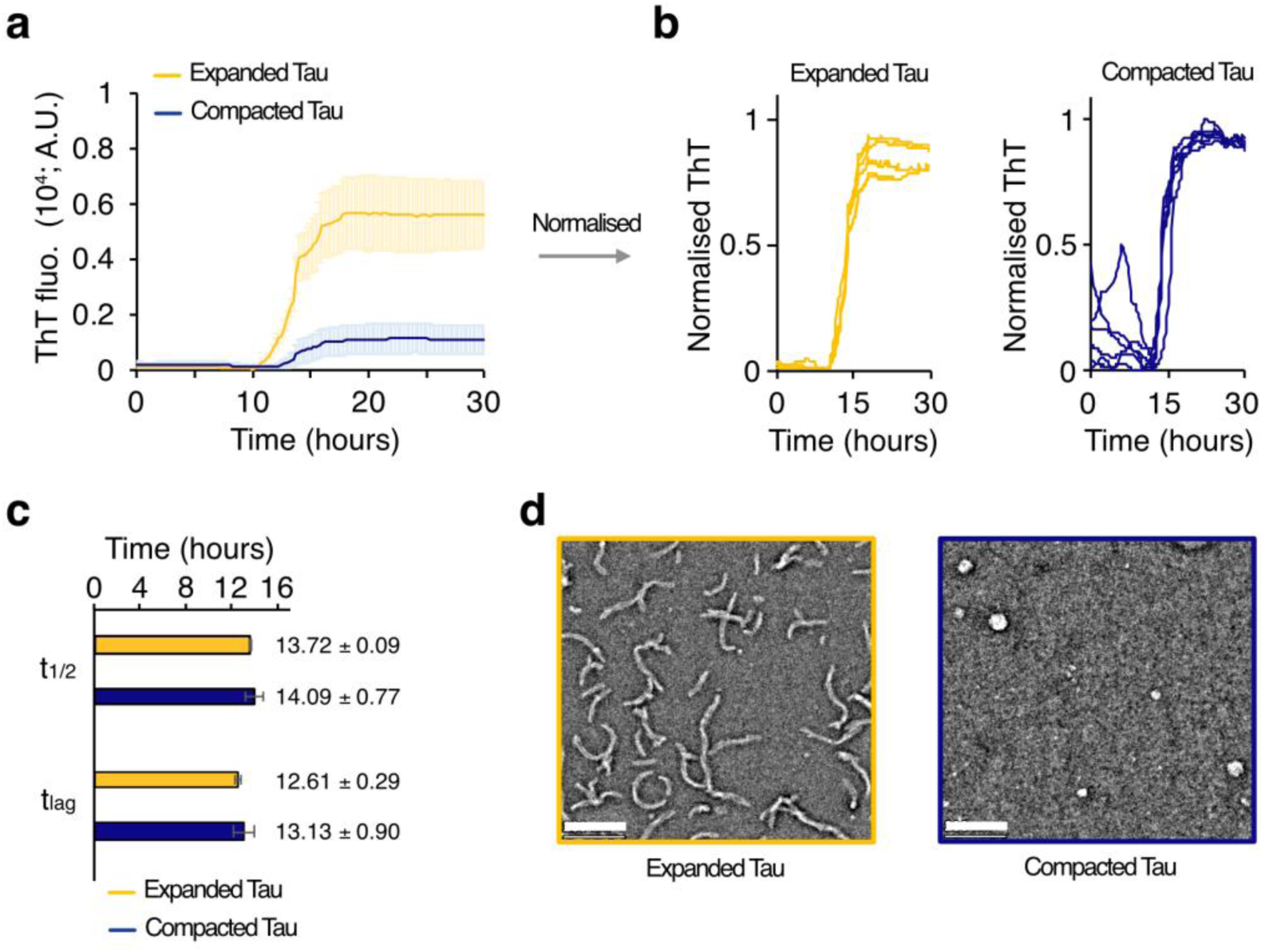
**a**, ThT aggregation kinetics of expanded (yellow) and compacted (dark blue) Tau, and **b**, their respective normalised ThT fluorescence, in 10 mM sodium phosphate, 5 mM TCEP pH 6 at 20 μM. **c**, The aggregation kinetics lag times (t_lag_) and half-times (t_1/2_) of expanded (yellow) and compacted (dark blue) Tau. **d**, Electron micrograph of expanded (yellow outline) and compacted (dark blue outline) at the end of aggregation kinetics. The white bar within micrographs indicates 200 nm.

## Discussion

In the present study, we demonstrate that monomeric Tau can populate two long-lived soluble conformational ensembles that exhibit markedly different self-assembly behaviours while remaining monomeric and intrinsically disordered in solution (Fig. 1a – f). Instead of representing a folded intermediate, the compacted state is best described as a dynamic ensemble stabilised by numerous weak intramolecular interactions, as demonstrated by the non-cooperative expansion of compacted Tau upon denaturant titration (Fig. 2d). In agreement with this model, our NMR analyses (Fig. 4a – c) identify segments with altered solvent accessibility containing several hydrophobic motifs, including aggregation-prone motif VQIVYK within R3 repeat and IVYK motif within R′ pseudorepeat as well as KxGS motifs that mediate microtubule binding^34–36^. Strikingly, it also includes regions known to bind Fyn kinase^37^ and muscarinic receptor binding sites^38^, which are involved in both physiological and pathological functions (Supplementary Fig. 6). Although our data do not implicate direct involvement of these motifs or residues for compaction (Fig. 3), they support a model in which weak, non-specific, fuzzy, multivalent interactions within the MTBR and its flanking regions collectively stabilise the alternative collapsed ensemble.

Our findings further suggest that the expanded Tau ensemble represents a metastable state that may follow two competing routes, intramolecular compaction and intermolecular phase separation, governed by the same underlying principles (i.e., electrostatic and hydrophobic interactions; Fig. 8). By reducing the ionic strength, the increasing Debye length could allow the negatively charged NTD and the positively charged MTBR to form intramolecular contact, creating a neutral environment. When the local charge within the MTBR is further neutralised under alkaline conditions, a regional collapse may take place and is likely stabilised by multivalent weak interactions, which may induce formation of denaturant-resistant structures as previously reported^22^, or entangled structures^39^. Meanwhile, above saturation concentration, the same physical principles can also favour intermolecular association. Rather than forming intramolecular contacts, the expanded ensemble may engage neighbouring molecules, promoting LLPS driven by the oppositely charged NTD and MTBR. In this scenario, condensation brings short-range interaction motifs into proximity, particularly KxGS and conserved hexapeptide motifs, which drive phase separation^40,41^, and stabilise the dense phase. Alternatively, Tau may also undergo fibril formation, which is mediated by aggregation-prone motifs VQIINK and VQIVYK within MTBR. Therefore, the balance of Tau stoichiometry and environmental conditions would determine Tau conformations and its phase behaviour.

**Figure 8.**
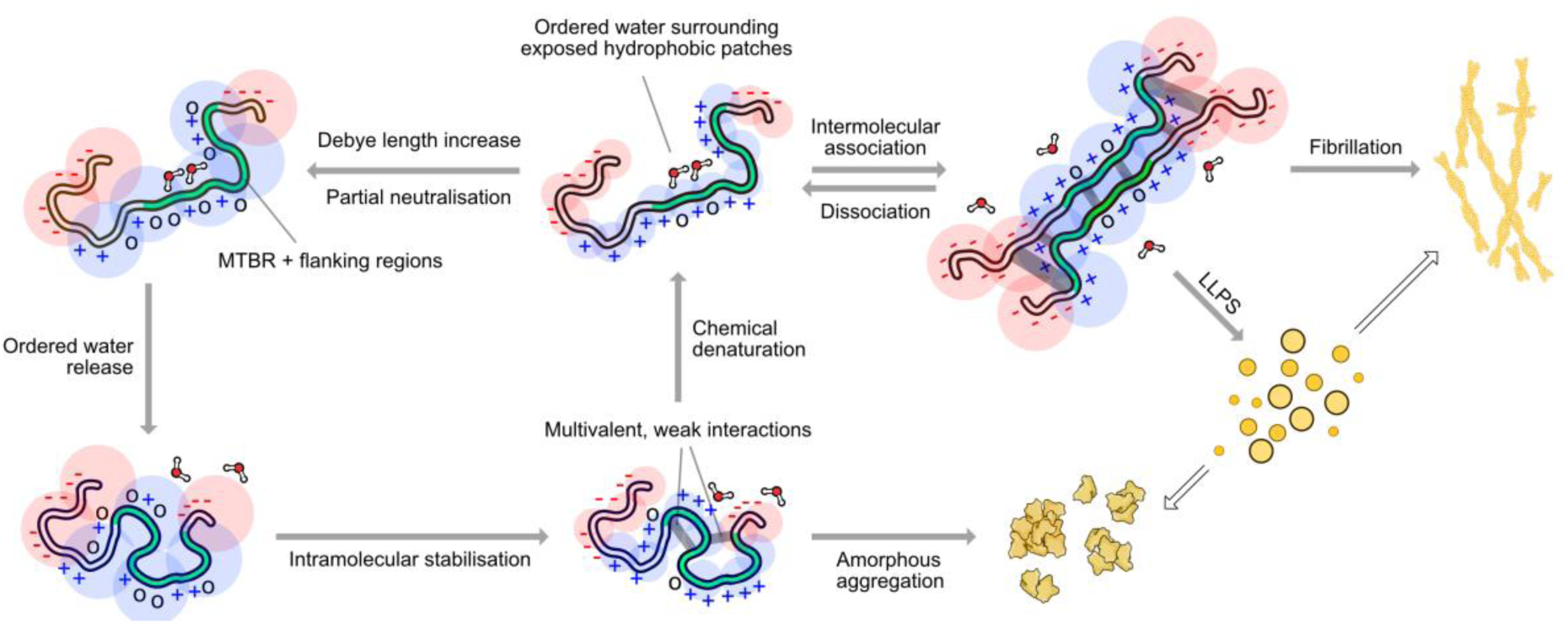
Cartoon model of charge-driven conformational change of Tau. Upon exposure to conditions that allow long-range intramolecular contact and partial neutralisation, expanded Tau samples numerous chain configurations. In this condition, chain rearrangement within the MTBR and its flanking regions releases water molecules surrounding hydrophobic segments and brings the chain closer together to form multivalent, weak interactions that further stabilise the collapsed state. In this state, Tau may re-expand upon adding denaturant to increase the solvent quality or diverge ‘off pathway’ forming amorphous aggregates. Alternatively, expanded Tau forms intermolecular contact with other Tau molecules, mediated by similar regions that collapse during compaction, which leads to the formation of liquid condensates or structured fibrils. Positive and negatively charge and its respective electrostatic field were visualised in blue and red, respectively. Grey arrows indicate proposed pathways based on results presented in this study, while white arrows indicate other possible pathways.

Collectively, our findings provide a mechanistic framework for understanding how electrostatic modulation can reshape the conformational landscape of soluble Tau and its subsequent self-assembly behaviour. In neurons, local PTMs, including serine and threonine phosphorylation, as well as lysine acetylation and ubiquitination, likely regulate these conformational biases by altering the electrostatic properties of the Tau chain^42–45^. Although these PTMs are thought to be coupled along the multi-step aggregation in disease^13^, our observations indicate that conformational remodelling and aggregation can be uncoupled. Therefore, we do not claim that the Tau species described here represent a direct pathological intermediate, nor that the proposed mechanism depicts a process in any disease. Instead, our results support a model in which a shift in the conformational energy landscape can be achieved solely by electrostatic perturbations, producing distinct conformational ensembles with different propensities for intermolecular association.

## Methods

### Expanded 2N4R Wild-type Tau expression and purification

The plasmid used for 2N4R wild-type Tau preparation, tau/pET29b, was a gift from Peter Klein (Addgene plasmid # 16316). The plasmid was transformed into Rosetta(DE3) competent cells (Novagen) for expression. The cells were grown in LB media with 50 ug/mL Kanamycin and 34 ug/mL Chloramphenicol at 37° C to an optical density of 0.6 – 0.9 at 600 nm before induction with 0.5 mM isopropyl beta-D-1-thigalactopyranoside (IPTG). For ^15^N labelled protein, bacteria were grown in isotope-enriched M9 minimal media containing 1g/L of ^15^N ammonium chloride (Sigma). The cells were harvested by centrifugation for 15 min at 10° C 14000 x g. The pellet was re-solubilised with lysis buffer (20 mM Tris, 500 mM NaCl, 2 M Urea and 5 mM Tris(2-carboxyethyl)phosphine (TCEP), cOmplete EDTA-free Protease inhibitor Cocktail (Roche), DNase I (Sigma), and RNase A (Sigma)). The pH of the buffer is adjusted to pH 8. Cells were lysed by sonication (3s on; 6s off at 50% amplitude for 5 mins). Cell lysates were centrifuged for 15 mins at 10° C 69000 x g. The supernatant was collected, titrated with acetic acid to reach pH 4, mixed gently until turbid, and neutralised with NaOH before centrifuged for 15 mins at 10° C 69000 x g. The supernatant was filtered and precipitated by adding 55% ammonium sulphate at 4° C for an hour. The precipitate was collected by centrifugation for 15 mins at 10° C 69000 x g and re-solubilised using lysis buffer. The re-solubilised precipitate was loaded onto gel filtration column HiLoad 16/600 Superdex 200 pg (GE healthcare) pre-equilibrated with lysis buffer using flow rate 0.75 ml/min. protein-containing fractions were analysed by SDS-PAGE, pooled and dialysed into buffer A (20 mM MES, 50 mM NaCl, 5 mM TCEP, pH 6) at 4° C overnight. The dialysate was loaded onto a HiTrap SP-HP column (GE Healthcare), pre-equilibrated with buffer A, and eluted with a 1 M sodium chloride gradient (buffer A + 1 M sodium chloride). Protein-containing fractions were pooled and dialysed into 10 mM sodium phosphate buffer pH 6 with 150 mM NaCl and 5 mM TCEP. The dialysate was aliquoted into low protein binding microcentrifuges tubes (Thermo Scientific), flash frozen in liquid nitrogen, and stored in -80° C. the purity of the protein was checked by SDS-PAGE and mass spectrometry. Protein concentration was determined using molar extinction coefficient e = 7450 M-1 cm-1 at 280 cm.

### Compacted 2N4R Wild-type Tau preparation

The purified expanded 2N4R wild-type Tau was diluted into <2 uM with 50 mM borate buffer pH 8-9 with 5 mM TCEP, dialysed using the same buffer at 4° C overnight and further dialysed into 10 mM sodium phosphate buffer pH 6 with 150 mM NaCl and 5 mM TCEP. The dialysate was separated from soluble oligomers and aggregates using Amicon Ultra 100k concentrator (Merck) followed by concentrating the flow-through using Amicon Ultra 10k concentrator. The concentrated flow-through, then, aliquoted into low protein binding microcentrifuges tubes (Thermo Scientific), flash frozen in liquid nitrogen, and stored in -80° C.

### Hydrodynamic radius determinations using microfluidic diffusional sizing (MDS)

Either expanded or compacted Tau were labelled with Alexa Fluor 488 C5 maleimide (Thermo Fisher) following the protocol provided. The protein and dye reaction was dialysed into buffer of choice to remove free dye at 4° C overnight. The dialysate was concentrated and aliquoted. The hydrodynamic radii were measured on a Fluidity One-M instrument (Fluidic Analytics) based on the machine operation principle^46^. Microfluidic circuits of the Fluidity One-M chip plate were primed using 4 μl of chosen buffer before adding 4 ul of sample (100 nM of fluorescently labelled Tau) to the chip. The Fluidity One-M instrument was operated according to the provided instruction.

### SDS-PAGE and Western Blotting

Samples were mixed with NuPAGE LDS Sample Buffer 4X (Thermo Fisher) 3:1 ratio and incubated on a heat block 90 °C for 10 minutes. Samples were then loaded and run in pre-cast NuPAGE 4 – 12 % Bis-Tris protein gels (Invitrogen) at 200 V for 30 minutes. Gels were stained with InstantBlue Coomassie Protein Stain (Abcam) for an hour and de-stained using water overnight to remove Coomassie background. For Western blotting, dry transfer was perform using the iBlot 2 Gel Transfer Device (Thermo Fisher). The unstained gel was stack into iBlot Transfer Stack (Thermo Fisher) according to the provided description. The transfer was performed using pre-programmed method (20 V for 1 minute; 23 V for 4 mintues; and 25 V for 2 minutes). The PVDF membrane rinsed with water and washed with Tris-buffered saline with Tween 20 (TBST) buffer (20 mM Tris pH 7.5, 150 mM NaCl, 0.1% Tween 20). The membrane washed blocked using blocking buffer (3% BSA in TBST) at room temperature for 1 hour. The membrane rinsed with TBST 5 times for 5 min. The membrane was incubated with Anti-Tau antibody (A98443; Antibodies.com) diluted into 1/1000 using blocking buffer for 1.5 hours at room temperature. The membrane rinsed with TBST 5 times for 5 min. The membrane incubated in the HRP-conjugated Goat Anti-Rabbit antibody (ab205718; Antibodies.com) diluted into 1/500 using blocking buffer for 1 hour at room temperature. The membrane rinsed with TBST 5 times for 5 min. The chemiluminescent ClarityWestern ECL Substrate mixed into 1:1 ratio with Clarity Western Luminol/Enhancer Reagent (Bio-Rad) and applied to the membrane. The membrane imaged with ChemiDoc Imagers (Bio-Rad).

### Far-UV circular Dichroism (CD) spectroscopy

Experiments were performed at 0.1 mg/ml over a 190-250 nm range at 25° C in 10 mM sodium phosphate pH 6.5, 5 mM TCEP and 150 mM sodium fluoride in 1 mm pathlength cuvettes. Spectra were acquired using an AVIV 410 spectrometer in 1 nm wavelength steps, averaged over three accumulations and baseline-corrected using buffer before smoothing, using the manufacturer’s software. Millidegree units were converted to mean residue ellipticity (MRE) with units deg cm^2^ dmol^-1^ res^-1^ using

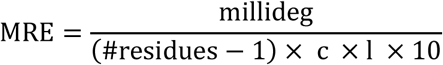

where c is the molar concentration and l is pathlength in cm.

### Fluorescence recovery after photobleaching (FRAP)

A Leica Stellaris 5 confocal microscope (Leica Biosystems) equipped with 63× oil immersion objective (Leica HC PL APO 63×/1.40 Oil CS2, NA 1.4) is used for imaging. Before imaging, sample was doped with 5% Alexa Fluor 488 labelled Tau. Sample was prepared by mixing the protein and polyethylene glycol (PEG) 8K to the final concentration of 20 μM and 10%, respectively. The sample was put on u-Slide VI 0.4 Bioinert (Ibidi). To perform FRAP, a 488 nm argon laser at 100% power was used to bleach a circular area 0.786 μm^2^ (radius = 0.5 μm) in a condensate with a diameter ∼ 2 μm. The recovery was measured by taking 1000 frames over 100 seconds (0.1 s per frame) after 0.3 s photobleaching. Curve were analysed and fitted using double exponential fitting to generate fitting values.

### Size exclusion chromatography-multiple angle light scattering (SEC-MALS)

The SEC-MALS system used was an AKTApure FPLC (GE), with an in-line Optilab T-Rex refractometer (Wyatt), and 8 detector Dawn Heleos II light scattering instrument (Wyatt). 500 μL of Tau sample (∼1 mg/mL) was injected onto a Superdex 200 Increase 10/300 GL column (GE Healthcare), pre-equilibrated in phosphate-saline buffer with 5 mM TCEP pH 7.2 and eluted at 0.5 mL/min directly into the on-line MALS instruments. Data was collected and processed using Astra 6 (Wyatt).

### Combinatorial droplet microfluidics and analysis

A custom-built microfluidic platform^33^ was employed for the combinatorial generation and imaging of water-in-oil droplets containing Tau and PEG 8K. Polydimethylsiloxan (PDMS)-based microfluidic devices were fabricated using standard soft lithography techniques as described previously^47^. The Tau and PEG 8K were pre-mixed with 5 μM Alexa Fluor 488 and Alexa Fluor 647 carboxylic acid (Thermo Fisher Scientific), respectively, to probe the concentration of the mixture. The temporal flow rates of Tau, PEG 8K, buffer and salt solutions as well as HFE-7500 oil containing 2% fluorosurfactant (RAN Biotechnologies) into the five different inlets were precisely modulated by a pressure-controlled pumps (OxyGEN, Fluigent). The oil phase flow rate was maintained at approximately 50 μL h⁻¹, while the total aqueous flow rate was set to 105 μL h⁻¹ forming water-in-oil droplets which incubated for ∼5 min while navigating microchannels prior to imaging using an epifluorescence microscope with excitation wavelengths of 488 nm and 640 nm.

Image analysis was performed following previously reported methods by filtering and classifying droplets that contain soluble or phase-separating moieties with a convolutional neural network. Concentration calibration curves generated by correlating intensity of the fluorescence probe with known concentrations of Tau and PEG 8K.

### Nuclear magnetic resonance

NMR experiments were performed on ^15^N-labelled expanded and compacted Tau with final concentration of 50 μM in 10 mM sodium phosphate pH 6.8 containing 10% D_2_O, 5 mM TCEP, 150 mM NaCl and 3 mM NaN_3_. The 2D ^1^H-^15^N heteronuclear single quantum coherence (HSQC), phase modulated CLEAN chemical exchange (CLEANEX-PM) and heteronuclear NOE (hetNOE) experiments were recorded at 293 K on Bruker DRX800 spectrometer (Bruker). Spectra were assigned by transferring full-length 2N4R Tau assignment^24^ and analysed using CcpNMR Analysis v. 2.4.2^48^. The chemical shift perturbation (CSP) comparing expanded and compacted Tau was determined using the following equation:

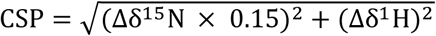

Heteronuclear NOE values were determined as previously described^49^ at 800 MHz. Briefly, interleaved 2D spectra were recorded in the presence and absence of a 4 s proton pre-saturation period. Peak intensities from the ‘on’ and ‘off’ spectra were compared to report on backbone dynamics on a picosecond timescale.

The solvent accessibility was probed by measuring the rates of chemical proton exchange (*k*_ex_) between backbone amide and solvent using CLEANEX-PM^31^. Fast HSQC spectra were acquired at five mixing times (4.92, 9.85, 19.7, 39.4, 78.8 ms) as well as a reference spectrum without mixing. The acquired data were fitted to obtain *k*_ex_ using the following equation:

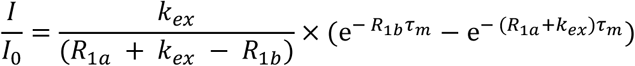

where I and I_0_ are the peak intensity at different mixing time τ_m_ and reference spectra, respectively; *k*_ex_ is the rate constant related to solvent exchange; R_1a_ is the combination of longitudinal and transverse relaxation rate in the CLEANEX-PM experiment, R_1b_ is the longitudinal relaxation rate for water (0.6 s^-1^).

### Dynamic light scattering (DLS)

The DLS experiments were performed on a Zetasizer Nano S instrument (Malvern Panalytical). Samples were prepared at various concentration and centrifuged for 10 mins at room temperature 20000 x g, with the exception of 200 μM expanded Tau, prior to transfer into BRAND^®^ UV cuvettes (Merck). Autocorrelation functions were recorded between 25 °C and 70 °C at 5 °C increment, with 5 minutes equilibration period at each temperature step. Below the onset temperature of liquid droplet formation, data were evaluated using distribution analysis resolving monomers and oligomers population present. Upon droplet formation, qualitative tracking was conducted to the apparent derived count rates (kcps) from Cumulants analysis algorithm. Both analyses were performed using the manufacturer’s software.

### ThT aggregation kinetics assay

Both expanded and compacted Tau were suspended at final concentration of 20 μM in 10 mM sodium phosphate pH 6 with 0.1 mM PMSF and 5 mM TCEP. Prior to shaking at 37 °C 300 rpm, ThT at a final concentration of 5 μM were added. ThT fluorescence was recorded at 15 minutes interval for 48 hours using PHERAstar plate reader (BMG Labtech).

### Transmission electron microscopy

After the end point of the aggregation kinetics assay, samples were collected and centrifuged at 10 °C 20000 x g for 10 mins. The sample, then, washed with 10 mM sodium phosphate buffer pH 6 with 150 mM NaCl and 5 mM TCEP and re-suspended with the same buffer to obtain 100 μM solutions. The concentrated sample applied to hydrophilic carbon-coated 400-mesh copper grids (EM Resolutions) and blotted the excess sample before stained with 2% (w/v) uranyl acetate. Fibrils were imaged using FEI Talos F200X G2 transmission electron microscope (Thermo Fisher Scientific).

## Acknowledgement

This work was supported by the European Research Council under the European Union’s Seventh Horizon 2020 research and innovation program, initially through the ERC grant DiProPhys (agreement ID 101001615, T.P.J.K.), and then through the award Research Studentship (Y.O.) by Wren-Cambridge Laboratory Project (RG8534_28778, T.P.J.K.).

We thank Aria Azari-Pour (Yusuf Hamied Dept. of Chemistry, University of Cambridge) for their assistance with the initial stages of protein purification; James London (Biophysics Facility, Dept. of Biochemistry, University of Cambridge) for their assistance with SEC-MALS; Heather Greer (Electron Microscopy Facility, Yusuf Hamied Dept. of Chemistry, University of Cambridge) for their assistance with TEM supported by EPSRC Underpinning Multi-User Equipment Call (EP/P030467/1); Dijana Matak-Vinković and Asha Boodhun (Mass Spectrometry Facility, Yusuf Hamied Dept. of Chemistry, University of Cambridge) for their assistance with LC-MS. We thank Sarah Perret and Si Wu (Institute of Biophysics, Chinese Academy of Sciences) for valuable discussions and feedback.

## Author contributions

Y.O., K.S. and T.P.J.K. designed and conceptualised the study. Y.O. and Z.H. performed the investigations. Y.O., Z.H., K.S. and T.P.J.K. provided the materials and methods. Y.O., K.S. and T.P.J.K. analysed the data. Y.O., K.S., and T.P.J.K. wrote the original draft of the paper.

