## Supplementary Figures for "Charge-driven remodelling of monomeric Tau determines its phase behaviour"

Yudisthira Oktaviandie *et al*.

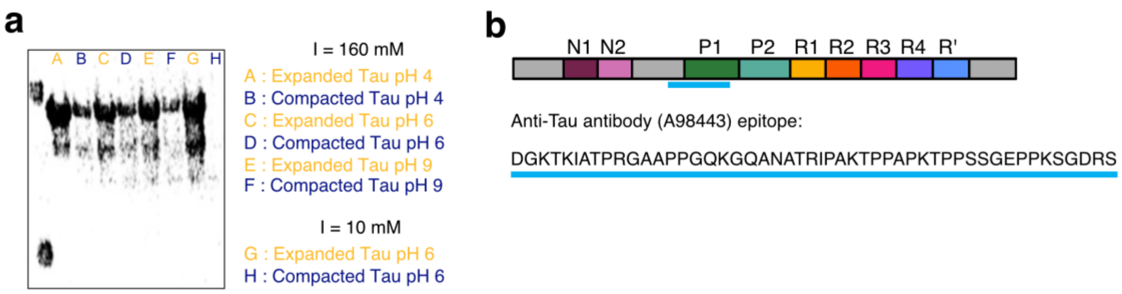


**Supplementary Fig 1**. **a**, Western Blot of expanded and compacted Tau previously incubated in different buffer condition. **b**, Anti-Tau antibody (A98443) epitope used in **a** and Fig. 1d within the main text.

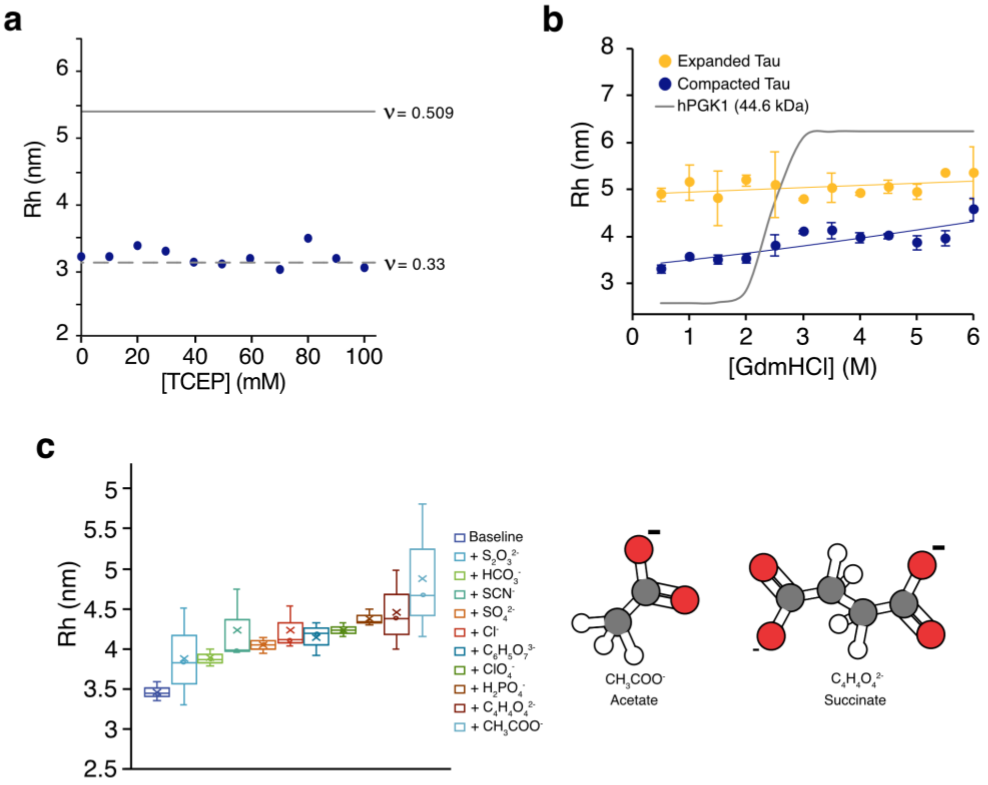


**Supplementary Fig 2**. **a**, Hydrodynamic radii of compacted Tau titrated with TCEP in 10 mM sodium phosphate pH 6. Different lines indicate expected molecular dimension as shown in Fig. 1c in the main text. **b,** Denaturant titration profile, and **e,** calculated $\Delta G_{expansion}$ of expanded (yellow) and compacted (dark blue) Tau compared with the folded human PGK1 protein (grey)[23] as comparison. **c**, Hydrodynamics radii of compacted Tau subjected to different Hofmeister salt at 200 mM. Depiction of the most potent salts for inducing Tau expansion, acetate and succinate, is shown.

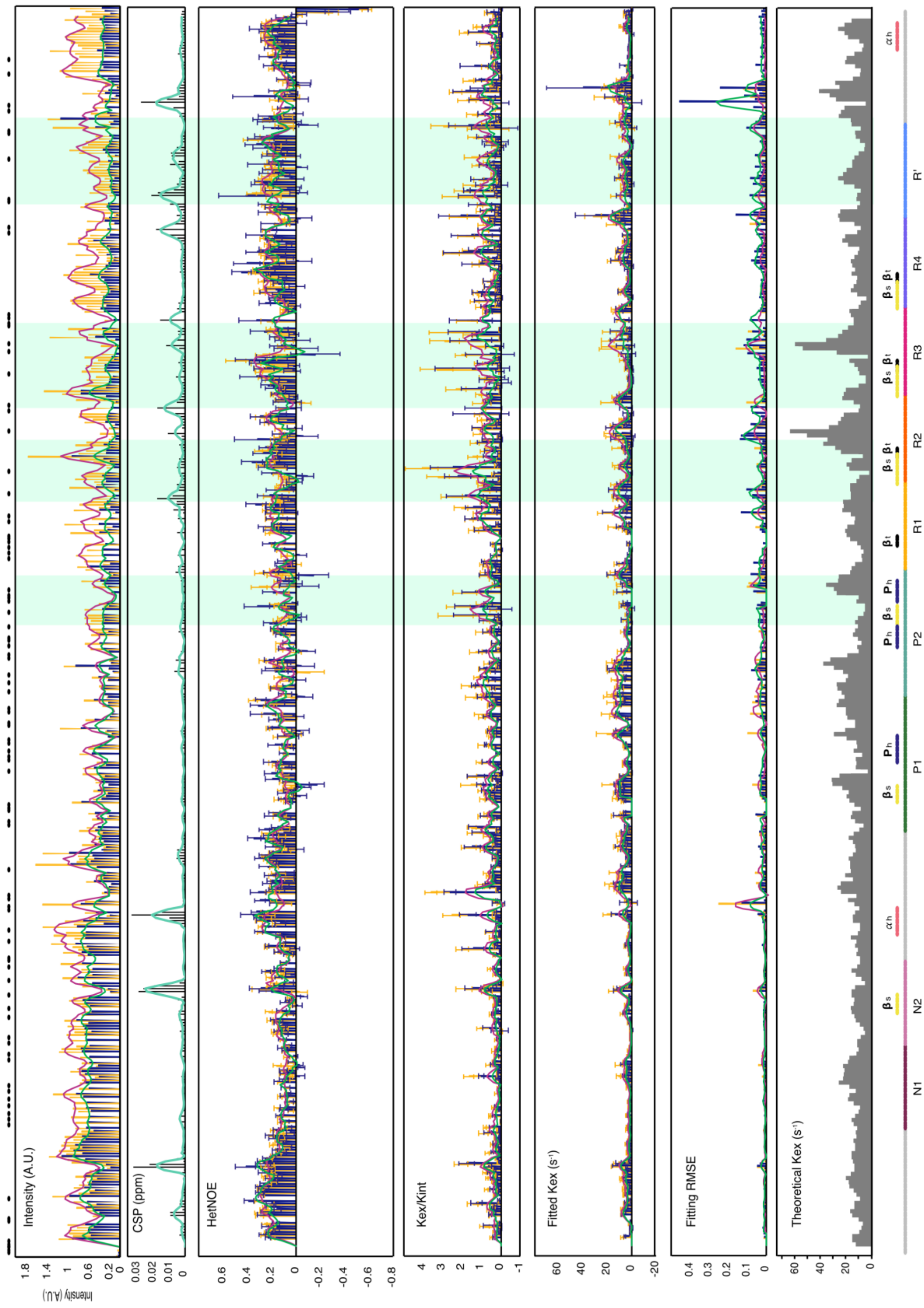


**Supplementary Fig. 3**. Intensity; chemical shift perturbation (CSP); HetNOE; $k_{ex}/k_{int}$ ratio; fitted chemical exchange rate ($k_{ex}$); fitting root-mean-square-error (RMSE); and theoretical $k_{ex}$, or $k_{int}$, of Tau. Expanded and compacted Tau are indicated in yellow and dark blue, respectively, with their respective interpolated trendline shown as magenta and green lines. Black dots at the top indicate unassigned residues. Coloured boxes at the bottom indicate different segments (N1 – R’) and transient structures ($\alpha_h$ = $\alpha$-helix; $\beta_s$ = $\beta$-sheet; $P_h$ = polyproline-helix II; $\beta_t$ = $\beta$-turn). The faded light green box indicates segments with significant solvent accessibility change shown in Fig. 2d within the main text.

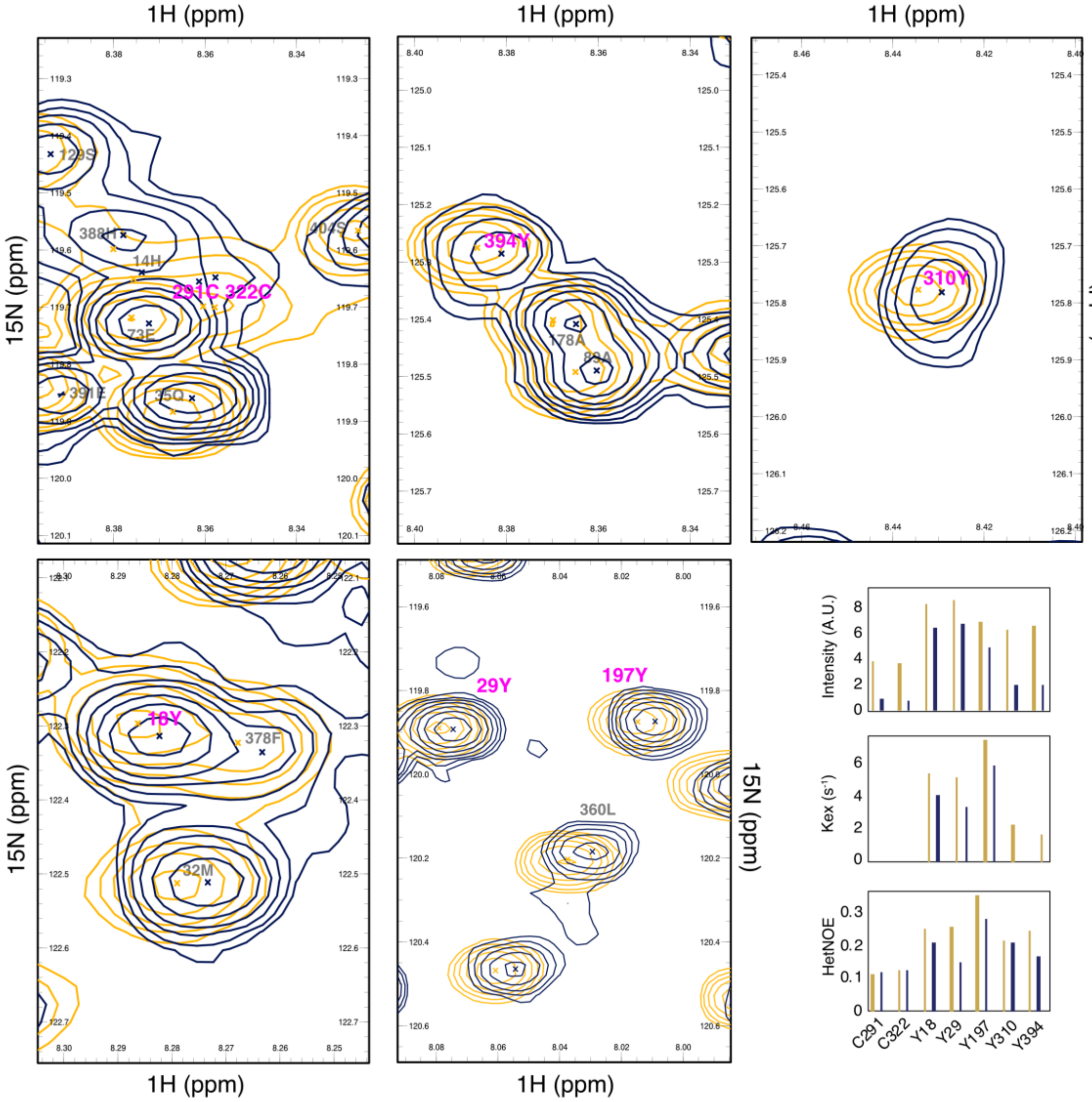


**Supplementary Fig. 4**. $^{1}$H-$^{15}$N-HSQC NMR spectra of expanded (yellow) and compacted (dark blue) Tau, as in Supplementary Fig. 2, indicating two cysteines (291C & 322C) and five tyrosines (18Y, 29Y, 197Y, 310Y, & 394Y) highlighted in magenta. Graphs at the bottom right correspond to the peak intensity, fitted $k_{ex}$ and hetNOE values of the cysteines and tyrosines.

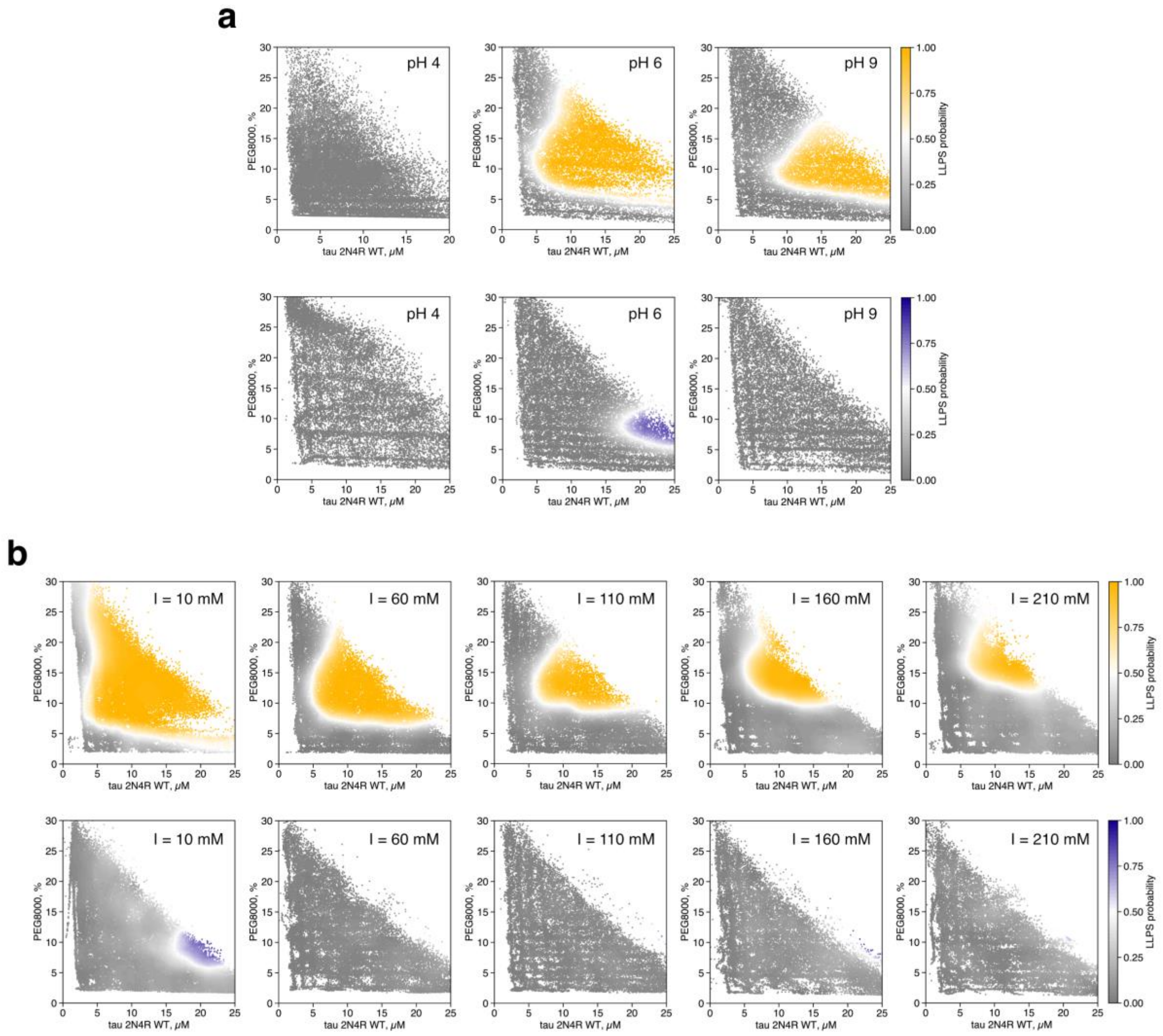


**Supplementary Fig. 5. a**, Phase diagrams of expanded (yellow) and compacted (dark blue) Tau in BR buffer at pH 4, 6 and 9, or **b**, in 10 mM sodium phosphate, 5 mM TCEP pH 6 with varying salt concentration.

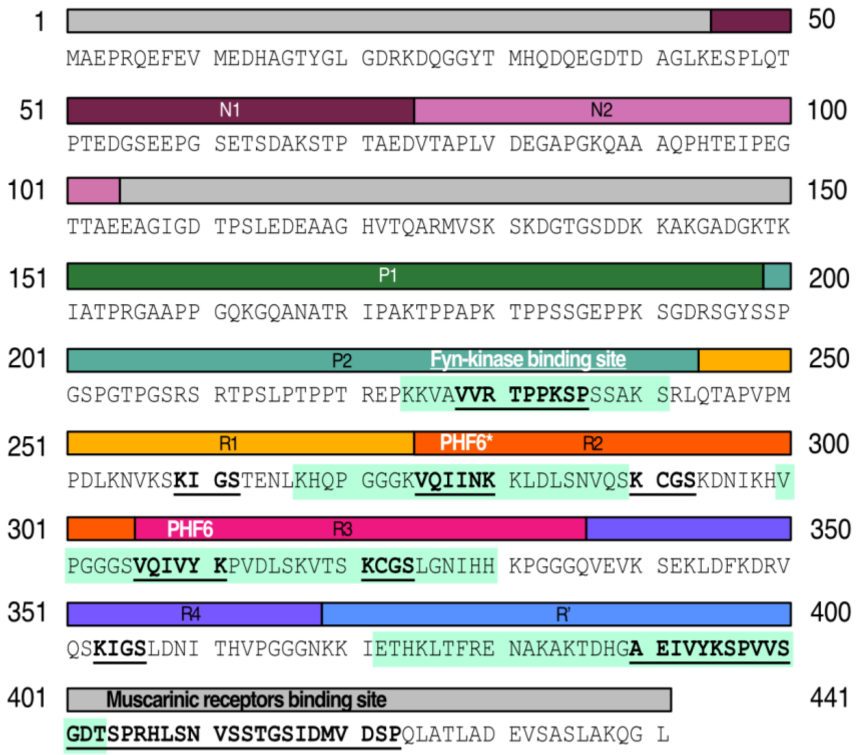


**Supplementary Fig. 6.** Full-length 2N4R Tau sequence highlighting known binding sites and regions implicated in compaction. The microtubule-binding region (MTBR) and adjacent flanking regions contain sequences implicated in Tau compaction (light green box). These regions overlap with previously identified binding sites, including those for Fyn kinase[37] and muscarinic receptors[37,38], as well as the aggregation-prone PHF6 and PHF6* motifs (bold and underlined). KxGS motifs, which contribute to microtubule binding and Tau phase separation[40,41], are also indicated.
